# Nonlinear brain connectivity from neurons to networks: quantification, sources and localization

**DOI:** 10.1101/2024.11.17.623635

**Authors:** Giulio Tani Raffaelli, Stanislav Jiříček, Jaroslav Hlinka

## Abstract

Connectivity is a widespread tool for the study of complex systems’ dynamics. Since the first studies in functional connectivity, Pearson’s correlation has been the primary tool to determine interdependencies in the activity at different brain locations. Over the years, concern over the information neglected by correlation has pushed toward using different measures accounting for non-linearity. However, one may pragmatically argue that, at the most common clinical observation scales, a linear description of the brain captures a vast majority of the information. Therefore, we measured the fraction of information disregarded using a linear description and which regions would be most affected. To assess how the spatial and temporal observation scale impacts the amount of non-linearity across multiple orders of magnitude, we considered fMRI, EEG, iEEG, and single-unit spikes. We observe that by treating the system as linear, the information loss is relatively mild for modalities with large temporal or spatial averaging (fMRI and EEG) and gains relevance on more fine descriptions of the activity (iEEG and single unit spikes). We conclude that Pearson’s correlation coefficient adequately describes pairwise interactions in time series from current recording techniques for most non-invasive human applications. At the same time, microscale (typically invasive) measurements might be a more suitable field for mining information on nonlinear interactions.

**Significance Statement:** In complex systems research, including neuroscience, the ubiquitous interest in network characterization by statistical dependencies (functional connectivity) invites increasingly sophisticated approaches. Various nonlinear measures, ultimately Mutual Information, emerge as alternatives to the conventional linear Pearson’s correlation coefficient. To fundamentally inform such decisions, we systematically assess the amount and reliability of non-linearity of brain functional connectivity across imaging modalities and spatial and temporal scales. We demonstrate more pronounced non-linearity in microscale recordings, while it is limited and unreliable in more accessible, non-invasive, large-scale modalities: functional magnetic resonance imaging and scalp electrophysiology. This result fundamentally supports the use of robust and easily interpretable linear tools in large-scale neuroimaging and brings essential insights concerning the non-linearity of microscale connectivity, including the link to brain state dynamics.

---

**M**any complex real-world systems, such as the human brain, Earth’s climate, or social or financial networks, do not easily allow their full intrinsic repertoire to be probed. Fully controlled laboratory experiments are often unattainable, and the researchers increasingly complement traditional theoretical or experimental approaches with advanced data analysis on observational recordings of their natural behaviour. A common starting step in such an endeavour is to capture the structure of statistical interactions between their constituent units. These interactions are then further analyzed, e.g., through graph theory, machine learning, or statistical approaches.

In the field of neuroscience, the term *functional connectivity* is used for such statistical dependence between remote neurophysiological events (1, 2). It has been the key cornerstone of the paradigm shift stressing the importance of understanding the mechanisms of spontaneous brain activity dynamics (3, 4). Over the decades, a plethora of functional connectivity indices has been proposed and applied. These indices form a densely populated zoo of methods, differentially sensitive to various forms of the concerned statistical dependence. This variety ranges from the classical Pearson’s correlation coefficient, assessing the strength of linear dependence between observed subsystems, to the principled use of Mutual Information. The latter is an entropy-based measure sensitive to any form of dependence, making it highly attractive for studying the interaction structures of highly nonlinear or even chaotic systems. Over the years, several papers have reported the effectiveness of MI on different modalities such as EEG (5) or fMRI (6) and significant changes in MI that correlate with disease (7, 8).

Mutual information as a measure of dependence may thus appear as a straightforward method of choice due to its lack of theoretical assumptions. However, in practice, some trade-offs need to be taken into account. The issue of estimate accuracy, the need for more extended time series (9), as well as computational demands, may favour more basic methods such as Pearson’s linear correlation coefficient. The pragmatic question the researchers (should) face is thus: Is the dependence pattern under study so exotic and nonlinear as to warrant using more general methods (or even, in principle, mutual information) at the expense of decreased statistical power, increased computational demands, and other costs of more advanced dependence quantification methods?

Many studies (10–13) have compared the quality of linear and nonlinear tools for specific tasks, mostly finding superior results with a linear approach. However, this always leaves a door open for a hypothetical yet another future approach to give better results in a given data situation. Here, we aim to establish more generally and quantitatively how much room is left for any potential future approach to improve over linear methods. To the extent that the linear part already covers the pairwise information, no approach can hope to yield significantly better results. To the contrary, more complex dependence estimators are prone to having a detrimental effect on the statistical power and subsequently on the sensitivity of the measures.

While the presence of non-linearity in brain signal dependence structure is theoretically well established and beyond dispute, much less is known about its strength and, therefore, relevance for further analysis, such as in functional connectivity networks. A principled approach for its quantification in terms of extra-Gaussian information has so far only been applied to a single modality (14). The investigation was limited to functional magnetic resonance imaging (fMRI) and at a single spatial signal averaging resolution given by the common, yet relatively coarse and arguably suboptimal Automatic Anatomical Labelling atlas (15). The results suggested that for this modality and at this spatial resolution, the nonlinear dependence contribution was likely negligible, supporting using Pearson’s linear correlation to quantify functional connectivity in this type of coarse fMRI data. However, this still left the question of the relevance of nonlinear dependences unanswered for a dominant proportion of neuroscientists, as the results could be only speculatively generalized to other neuroimaging modalities and spatial scales.

To at least partially fill this gap, we here apply the principled methodology for non-linearity quantification to neuroimaging data ranging across modalities from fMRI through scalp electroencephalography, intracranial electroencephalography, to single neuron (spike) recordings, adding analysis across orders of magnitudes of spatial averaging, and additional study of the sources, reliability, and localization of the observable non-linearity. Moreover, we provide an implementation in open code, allowing to quantify non-linearity in other datasets in neuroscience and beyond, which should provide a powerful tool also to researchers in different disciplines, following early successes of similar methodology in climatology (16) and financial networks (17).

## Results

Probably the simplest, and in some sense canonical, form of dependence between quantitative variables is that of a purely linear relation between two variables in a jointly normal (Gaussian) distribution, the strength of which is captured by the well-known Pearson’s (linear) correlation coefficient. Indeed, linear correlation is even sufficient to determine mutual information in a bivariate Gaussian distribution through the simple equation (18):

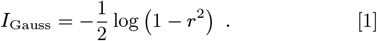

It is precisely the deviation from the jointly Gaussian distribution that makes the use of Pearson’s linear correlation coefficient potentially suboptimal, calling for more advanced methods such as the universally sensitive mutual information. The strength of the deviation from a Gaussian dependence pattern is thus the object of our interest. Moreover, we are not concerned with the non-Gaussianity of the marginal distributions individually (as these can be easily treated by using monotonic nonlinear rescaling such as log-transform or application of Spearman’s correlation instead of Pearson’s). We look particularly at such deviations from Gaussianity that concern the copula. This captures the relation between the variables and entails the full characterisation of the dependence structure, being invariant with respect to any bijective monotonic rescaling of marginals.

Note that, to avoid too technical language, we shall use the terms “linearity” and “Gaussianity” (of distribution/dependence/interaction/pattern/process) interchangeably throughout this manuscript. However, in a strict sense, linearity is a bit wider term in particular contexts (e.g., imagine linear functional dependence as the best fit between two variables with non-Gaussian distributions of errors or a linearly coupled multivariate process with nonlinear driving noise).

We shall leverage the well-known statistical physics result that the bivariate Gaussian has the maximum entropy, i.e., the minimal information (under fixed, e.g., Gaussian, marginals), among the distributions with the same correlation. Thus, every joint distribution that doesn’t have a Gaussian copula (we shall call these distributions “nonlinear” ) has higher MI than expected from correlation by the formula Eq. (1). This property allows us to define the distribution non-Gaussianity by

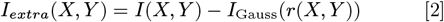

To stress that *I*(*X, Y* ) contains both linear and nonlinear contributions, we will later refer to this as Total Mutual Information (TMI), which we by default estimate with a binning method (see “MI estimator” in the methods section and the SI for alternatives). To get a fair comparison and better control of estimator bias, instead of using *I*_*Gauss*_ from Eq. (1), we use the Gaussian Mutual Information (GMI) estimate provided by applying the same estimator to linear surrogates of given data.

For illustrative purposes, we will use a normalised variant of *I*_*extra*_ that we call Relative Non Linearity (RNL), defined as:

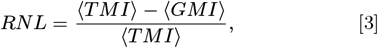

where the average ⟨·⟩ is over all pairs of time series. To give a sense of what different values of RNL imply, we show in Fig. 1 four examples sampled from bivariate distributions.

**Fig. 1.**
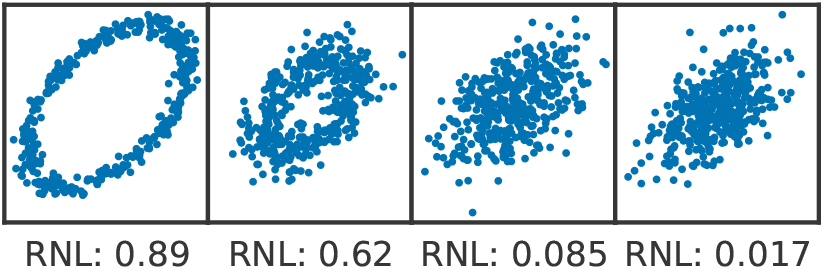
Values of RNL for four samples from bivariate distributions with different nonlinear content. The rightmost sample is from a bivariate normal distribution with *ρ* = 0.5.

We identify two primary sources of apparent non-linearity in time series observations: intrinsic non-linearity and non-stationarities. Intrinsic non-linearity happens when the recorded samples are genuinely identically distributed but do not have a Gaussian joint probability distribution. For example, this can include a relationship between absolute values or out-of-phase synchronization phenomena.

We refer to non-stationarity when the samples are not identically distributed, but the source distribution depends on time. The samples will then, generically, not be distributed according to a multivariate Gaussian, even if the source distributions are all Gaussians. Most estimators require the samples to be i.i.d. or at least from a stationary distribution. On the other side, the non-stationarity, even of resting-state (rs) data, is well known and studied on its own as a potential source of insight into the brain functioning (19, 20). The effect’s magnitude depends on the estimator and the non-stationarity’s properties. Examples of non-stationarities are the switching between states—each associated with a different correlation, isolated bursts of activity, or a continuous drift of mean, variance, or correlation.

Along with these truly neural sources for the observed non-linearity, others can reside in the acquisition and pre-processing of the signal. For instance, non-monotonous transformations may easily result in an increase in observed non-linearity, and in particular, observation of non-linearity between originally linearly related processes (it is illustrative to imagine the effect of taking, as a domain-specific preprocessing step, absolute value or square of Gaussian signals before probing their functional connectivity; a similar but less straightforward effect is obtained by working on, e.g., Hilbert-transformed data or (band-limited) signal power time series, a transformation much more common in neuroscience).

### Strength of non-linearity

We begin by measuring the fraction of information in all connections not explained by a bivariate Gaussian copula. We introduced this measure as Relative Non-Linearity.

We start from rs-fMRI and look at RNL over a range of region numbers and sizes to probe the effect of different degrees of spatial averaging. We observe a consistent presence of non-linearity for the different region sizes from the Craddock atlas (21). The fraction of MI not explained by the correlation (Fig. 2A) sits around 4% for all region sizes except for very few and large regions. At the same time, the non-linearity observed in the shadow (phase randomized) dataset remains below 2%, providing a measure of the estimator bias in the absence of non-linearity (this shadow dataset is used to control for this minor overestimation of observed non-linearity from finite-sized samples by our methodology). Note that, albeit quantitatively minor, the difference in MI between empirical and shadow datasets is always statistically significant, except for the atlas with ten regions (when applying a strict Bonferroni correction for multiple comparisons).

**Fig. 2.**
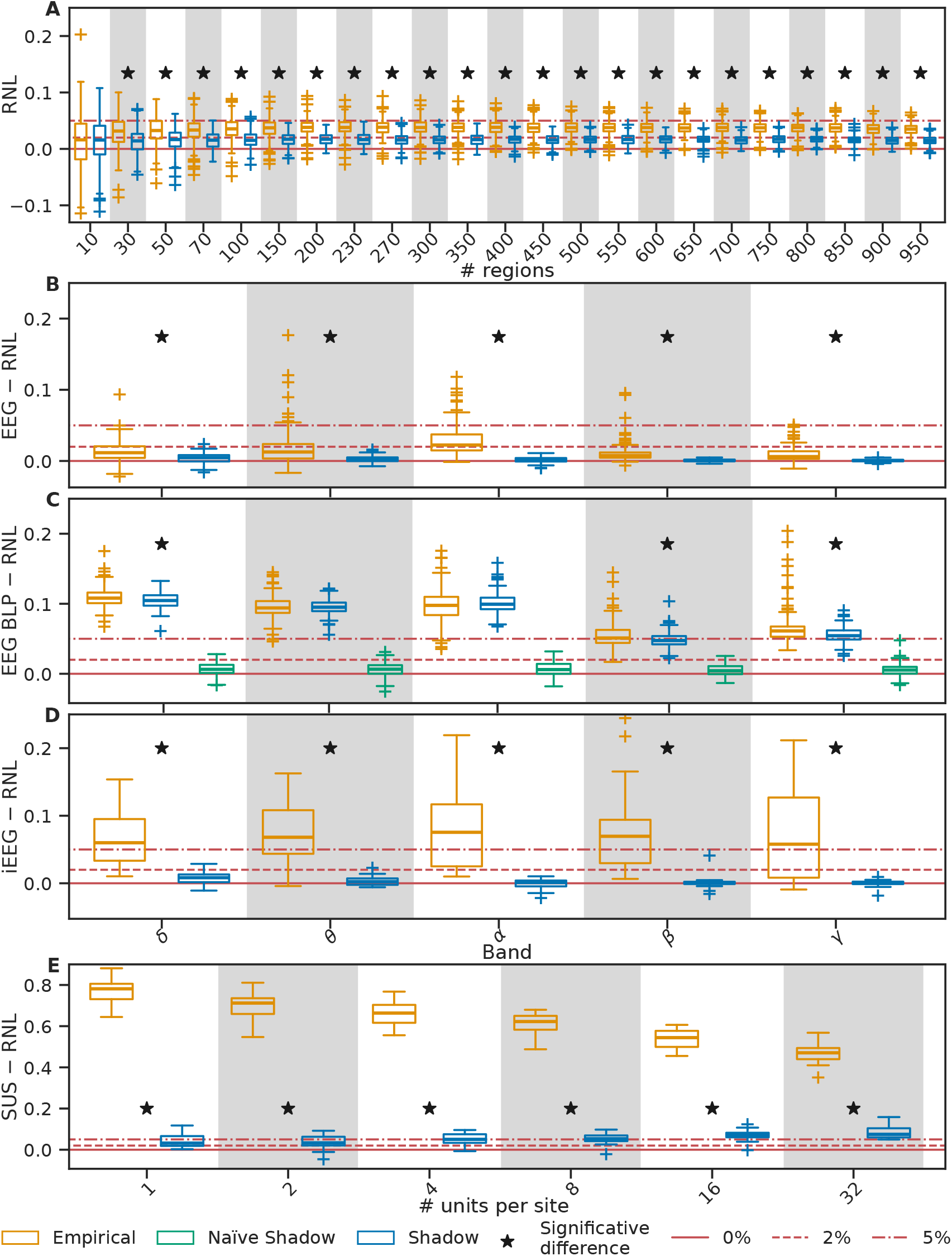
Distribution over subjects of the Relative amount of Non-Linearity (RNL). A) fMRI data using Craddock parcellation and a varying number of regions, B) EEG scalp voltage, C) EEG band-limited power, D) iEEG voltage, E) Single Unit Spikes instantaneous firing rate. The shadow dataset is a linear surrogate of the empirical one; see text for details. The black asterisk marks a significant difference between the Empirical and Shadow datasets. The red lines mark values of 0%, 2% and 5% RNL as a reference for the eye.

In the EEG time series, we observe a significantly higher presence of non-linearity (*p <* 0.05, one-sided t-test, Bonferroni corrected) than in the shadow dataset for all frequency bands. However, the RNL in EEG tends to be even lower than in fMRI, particularly under 2% (Fig. 2B), except for the *α* band. The variable number of samples available in each band explains the dependence between the frequency band and RNL in the shadow dataset. See the Methods section and the SI for details.

In iEEG, non-linearity is more prominent than in the previous two datasets. For all bands, the empirical RNL tends to be above 5%. At the same time, in the shadow dataset, it is confined below 1% (Fig. 2D). However, nonstationarities due to interictal epileptic activity explain part of the non-linearity (see SI). Moreover, the iEEG was acquired during natural activity (neither standard resting state nor sustained task), which might affect the stationarity of the sequences and their RNL.

Lastly, we looked at the RNL in single-unit spike rates in mice. The relative contribution of non-linearity is above 40% (Fig. 2E) in all cases, over the full range of group size averaging, between 1 and 32 units. We observe the highest values of RNL for the smallest group sizes, and averaging over larger groups reduces the RNL. We found no correlation between the RNL and the number of regions in a given mouse (ranging between 9 and 16). Thus, we present the results across all mice.

At the same time, the RNL never surpasses 10% in the shadow dataset. The magnitude of the bias increases for the average of larger groups, as does the autocorrelation in the series.

### Beyond mutual information

Mutual Information guarantees that any non-linearity emerges as a positive difference compared to linear surrogates. Other measures of connectivity do not offer equally general and interpretable quantification of nonlinearity. However, in their specific scenarios, they may be in principle more sensitive; moreover, if the measures include multiple time points or state-space representation (as is common for directed or effective connectivity measures), one can never entirely disprove the existence of some complex nonlinear pattern invisible in a joint bivariate marginal.

Thus, although previous studies have reported surprisingly good performance of linear approaches across modalities (10, 22), in line with our observation of negligible nonlinearity, to further reasonably explore and verify that the reduced role of non-linearity is not due to the specific measure we adopted, we tested selected alternative measures of connectivity for possible better sensitivity compared to MI. In the presence of nonlinear distributions, the connectivity matrices (obtained with any measure) from the empirical data should differ from surrogates more than the connectivity estimated on linearised, shadow data.

We estimate the sensitivity of a measure as the quality of the separation of empirical and shadow data based on the average difference from surrogates of the connectivity matrices. In particular, we computed connectivity matrices with Effective Connectivity estimated via Conditional MI KNN (using Tigramite (23)), Chatterjee correlation (24), and distance-transformed Chatterjee correlation (25) with different values of lag for the embedding. For EC, we observe a separation between empirical and shadow similar to the one with MI. This suggests that EC, like FC, can be well approximated with linear models, as done for example in (12). We observe lower separation for the approaches based on the Chatterjee correlation, in some cases, close to chance.

Our analysis suggests that the non-linearity is already well captured with information-based FC; see the SI for more details.

### Localization

As described above, non-linearity is relatively small for the more accessible non-invasive neuroimaging modalities. We conjecture that high temporal or spatial averaging in fMRI and EEG masks most of the non-linearity. However, it might still be consequential if localized in specific regions. For instance, the authors in (26) suggest localization of non-linearity in the occipital region. We thus further evaluated non-linearity’s localization, looking at regions participating in consistently nonlinear connections across subjects.

For fMRI data, we used the AAL90 atlas as it provides explicit anatomical labelling for the regions where the non-linearity might be localized. Under stringent preprocessing, the localization of nonlinearity was mostly concentrated to connections within the occipital cortex and was relatively weak (of similar strength as in the shadow datasets), and in particular held significant similarity (correlation between region degrees 0.37, *p* = .00028 uncorrected) between the spatial localization of non-linearity in the empirical and shadow datasets (see methods and SI). Notably, the localization of the non-linearity and the similarity of its spatial pattern between the empirical and shadow datasets were gradually more apparent with reduced denoising steps in preprocessing (moderate preprocessing, or raw data variant). This trend suggests that most region-specific non-linearity is due to artefacts and is removed during preprocessing.

EEG data (Fig. 3) offer a different picture. Here, substantially nonlinear relationships are pretty limited in the *θ* band (only 38% channel pairs) while present in more than 95% of channel pairs in other bands. In the shadow dataset, less than 0.8% of the relationships are consistently nonlinear across subjects.

**Fig. 3.**
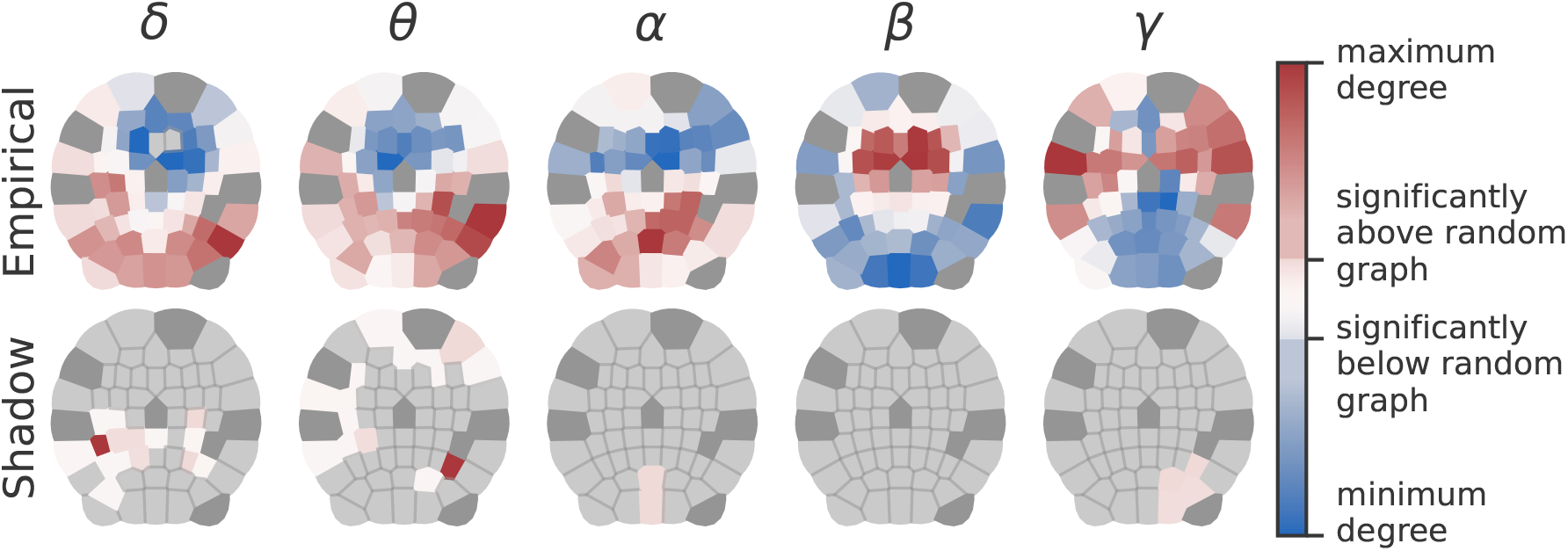
Degree of the regions in a network weighted by the z-score of TMI compared to surrogates. Light grey regions have degree zero. Dark grey regions have been excluded from the analysis as they are often missing or corrupted. The representation is a Voronoi tessellation of the stereographic projection on the xy-plane of standard electrode positions. The nasion is facing up.

Overall, in the EEG data, the non-linearity is spatially heterogeneous. For the slower bands *δ, θ*, and *α*, we observe that the degree of the occipital regions is higher (and in frontal regions lower) than expected from a random graph, suggesting localization of non-linearity. Conversely, for the high-frequency bands *β* and *γ*, the results show a predominance of non-linearity in frontal and temporal electrodes. This anterior-posterior gradient may be partially attributed to the spatial distribution of the most prevalent sources in normal awake EEG (27), yet more research is warranted.

### Reliability

Despite relatively small non-linearity, considering it may still be beneficial if the additional information is sufficiently stable over repeated measures. Therefore, as the last test, we looked into the non-linearity reliability across sessions. For each region pair, we measured it as the Spearman’s correlation of the subjects’ connectivity in different sessions. We compared the connectivity based on Total Mutual Information (TMI, i.e., accounting for non-linearity), Pearson’s correlation, and *I*_Gauss_ applying Eq. (1).

We observe that the contribution of non-linearity is limited and unreliable under our measure. The reliability of Pearson’s correlation is, on average, across connections, more than twice that of the reliability of TMI (0.277 against 0.134). The reliability of the *I*_Gauss_ (which is equivalent to the reliability of the absolute value of interregional correlation) is similarly low as TMI reliability (0.153). This suggests the advantage of keeping the sign of the interaction.

Moreover, *I*_Gauss_ is not only comparably reliable as TMI, but we observed that it predicts TMI in a different session even slightly better (0.008 or 5% increase in Spearman’s correlation) than TMI predicts itself. This shows the practical advantage of using a linear functional connectivity measure at this scale.

See SI for detailed results and analogous analysis on EEG and iEEG data, which offers a similar picture, excluding any clear advantage of TMI.

### Effect on classification

While at first sight the choice of the method is indifferent in case of (almost) linear dependencies, choosing nonlinear methods can come at a practical cost. This can be in terms of computational demands or lower statistical power, as mentioned in the introduction, or as sensitivity to additional noise, as suggested while assessing the reliability.

To elucidate the practical (potentially detrimental) effects of applying estimators including nonlinear contributions, we repeated part of the analysis in (11), including MI measures.

In particular, we trained a logistic regression classifier (the best-performing classifier in the benchmark) to distinguish schizophrenia subjects and controls. The features were TMI, GMI (surrogates after marginal normalisation), MI in shadow (surrogates before marginal normalisation), Pearson’s correlation (before and after marginal normalisation), or Spearman’s correlation.

While the results for the different features are pretty similar, the only significant difference is the improvement when using GMI instead of TMI. See more in the SI.

### Sources

To better understand the relevance of observed non-linearity, we will now survey some sources of non-linearity that may be considered spurious.

EEG offers an example of non-linearity derived from the choice of the observable. Let us consider band-limited power and compute the shadow dataset naïvely from the sequence of power values (modulus of the Hilbert transform). This would be an acceptable choice as, for example, the computation of FC from the signal envelope is customary (28, 29) in MEG. We observe in all bands a 5% to 10% share of the information not described by the linear dependence (Fig. 2C, yellow boxplots). However, we also observed comparable non-linearity when extracting the power from a surrogate of the original EEG time series, documenting that most non-linearity arises from power computation (Fig. 2C, blue boxplots).

In most bands, the non-linearity in the shadow dataset is still lower than that of the empirical one. This suggests that the transformation from voltage to power is not responsible for the entirety of the observed non-linearity in bandpower dependence. However, the large amounts of spurious non-linearity mask the significance of band *α* (Fig. 2B).

As a potent source of apparent non-linearity, as mentioned in the introduction and discussed in detail for climate systems elsewhere (16), one should pay attention to non-stationarities. An obvious potential source of non-linearity in the iEEG data is epileptic activity. We observe how (Fig. 4), measuring the RNL on a sliding window across the recording of a seizure, up to almost 90% of the total information appears to be due to non-linearity.

**Fig. 4.**
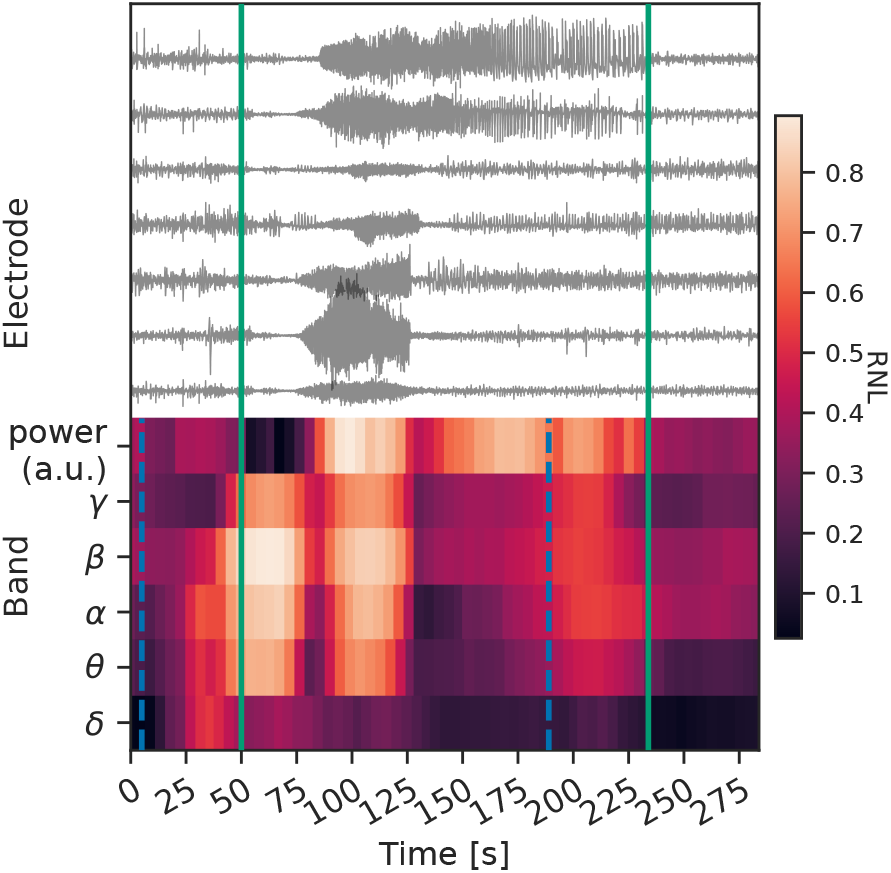
Sample seizure showcasing non-stationarities’ effect on the apparent non-linearity. Top: traces from a subset of electrodes. Bottom: mowing window values of RNL and band-limited power. The green solid lines mark the beginning and the end of the seizure. The blue dashed lines mark the beginning of the first windows containing samples past the green lines.

In particular, we observe how the RNL is sensitive to the signal’s amplitude variation. The first peak for bands *θ* to *γ* happens when the sliding window crosses between the initial regime of low power and the first high amplitude oscillations. The RNL decreases when the window contains only wide oscillations and increases again when the first group of electrodes reverts to lower power. The last increase in RNL happens when the windows cross out of the seizure.

A second example of the effect of non-stationarities comes from spiking data of individual neurons. Over the 30 minutes of recording, even without stimuli, the mice’s brain activity had the opportunity to drift or switch through different states (most notably, immobile and running periods). Computing the RNL over shorter windows with fewer state changes and then averaging, we observe a larger fraction of the information described by the linear structure and thus a reduction of the non-linearity, see Fig. 5. However, even in the last case of 3-minute epochs, the average RNL stays much above what is observed with other modalities.

**Fig. 5.**
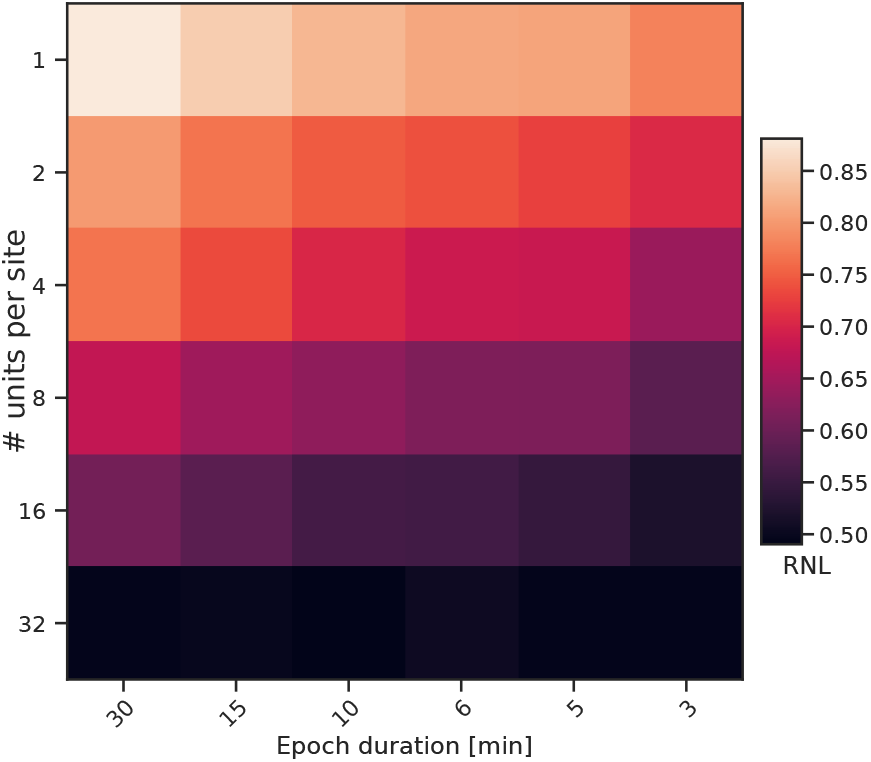
Average RNL across mice varying the number of units and the epoch length.

## Discussion

The fundamental non-linearity of neuronal activity leads to a relevant fraction of nonlinear information in single-unit spikes. However, moving to more accessible and non-invasive human data, the values are vastly reduced. We conjecture that this might be an effect of the scale at which the activity is recorded. Indeed, fMRI and EEG imply large spatial or temporal averages. At every step of the progression from groups of units to iEEG to EEG, the number of neurons involved grows by a factor ≈ 10^4^. The Central Limit Theorem alone could explain much of the linearising behaviour. Demonstrating this intuition using fully realistic models presents a considerable challenge. Nevertheless, as shown in the Supplementary Information, already a classical system of Kuramoto oscillators can illustrate a gradual decay in observable RNL across three orders of magnitude of spatial averaging, closely paralleling the relationship observed between iEEG and EEG in the most nonlinear subjects. For other possible mechanisms suppressing non-linearity, see (13).

The more significant fraction of nonlinear information in the iEEG dataset might support this interpretation. Part of this non-linearity may be explained by non-stationarities in the form of IEDs affecting the higher frequency bands and drift or state switching due to the natural activity of the subjects during the recording. The surge in the values of RNL in correspondence with the variations in amplitude during seizure supports the interpretation of the bursts in high-frequency during IEDs (non-stationarities) as sources of observed non-linearity. The apparent contrast with the results in (13) might be the result of a different sensitivity to non-stationarities of our approach or to the presence of significant unexplained variance in all models in (13), which could be exploited by a future (non-) linear approach to improve predictions. However, this modality remains more promising for detecting nonlinear interactions in the human brain, and future work will have to test this with improved control over non-stationarities.

While the presence of some non-linearity is significant at all scales, its variability at the individual level makes it hard to assess its per-subject localization, requiring group-level analysis. Indeed, the non-linearity is reduced and unreliable across sessions, which could be explained by a substantial contribution of noise in the difference between the Gaussian and total MI. While we have observed spatial heterogeneity of the observed non-linearity in both fMRI and EEG, it is technically nontrivial to immediately discern to what extent this (spatial pattern of) observed non-linearity is due to spatial heterogeneity of artifacts, nonstationarities, or genuine nongaussian dependence patterns. Elucidating this requires further study, tackling sampling issues for studying nonstationarities across timescales, inter-subject heterogeneity, as well as disentangling the plethora of possible artifacts at play at each modality.

Non-linearity is present at all levels of our analysis, and the oblivious use of Pearson’s correlation might be inappropriate. However, we advocate caution when designing studies that plan to apply MI or other nonlinear connectivity measures. Without accurate control of spurious sources of non-linearity, the improvement from using MI instead of correlation might be hardly reproducible, if any, and far outweighed by the technical complexities of nonlinear measure applications, including decreased statistical or classification power. Similarly, in fMRI, the minor nonlinearity has already been shown to have a negligible effect on graph theoretical measures (30). We expect similar effects for other modalities with negligible nonlinearities; however in case of crucial (e.g. clinical) applications a targeted application of nonlinear measures may be warranted, in line with the view that under certain (typically pathological, such as epileptic seizure) circumstances the normally very high-dimensional (apparently linear) brain dynamics collapses into low dimensional oscillatory (nonlinear) mode (31). For this reason, we advocate for using linear tools unless specific analysis displays a significant presence of nonlinear behaviours and excludes any spurious sources.

Lastly, many FC studies seek dependencies in the power or frequency domain, particularly in EEG (32), where there is an established relationship between frequency and function. We recommend careful treatment of power-based connectivity, as its computation itself introduces nonlinear effects. Concerning frequency-resolved analysis, while we have observed statistically significant, yet quantitatively minor nonlinearity using our methodology, there is an important debate around the need to go beyond Gaussian dependencies with approaches looking at higher-order spectral properties showing the signature of non-linearity (33, 34). Future work will have to elucidate the relation of particular definitions of non-linearity and higher order effects adopted, and to quantitatively (beyond pure statistical testing of the binary question of presence or absence) assess the amount of non-linearity and the role of averaging in the frequency domain, to provide a unified perspective. Biologically realistic simulations of neural networks, and linking to patterns observed in multimodal datasets such as concurrent EEG-fMRI (35, 36), may play a key role in shaping our understanding of the observed (non)-linearity across spatial, temporal, and frequency scales.

## Materials and Methods

To estimate the nonlinear content in the relationships between regions, electrodes, and units, we compared the Total Mutual Information (TMI) between time series to the estimate from surrogates where only the linear relationships are preserved. This allows us to evaluate the global amount of non-linearity and which in regions or electrodes it is more substantial.

### MI estimator

We estimate the MI of two variables through equiquantal binning (also known as equiprobable (37)): the samples are sorted on a grid with the same bin number for both variables. The bins for each variable have variable width so that the sum of the counts of the bins over the other variable always gives the same number of data points. The MI is then computed from the estimated probabilities of each bin *p*_*ij*_ as the difference between the sum of the marginal entropies of the two variables and their joint entropy:

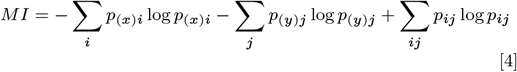

with *p*_(*x*)*i*_ = ∑_*j*_ *p*_*ij*_, *p*_(*y*)*j*_ = ∑_*i*_ *p*_*ij*_, and *p*_(*a*)*l*_ ≃ *p*_(*a*)*k*_, *a* = *x, y*, ∀ *l, k* ∈ [1, *N* ] where the equality holds for all *l* and *k* only if the number of bins *N* is a divisor of the sequence length *S*. We chose 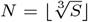 in line with the previously recommended pragmatic heuristic (38).

This estimator (as MI estimators in general) is known to be affected by biases. Here, in particular, for high values of MI, the estimate is bounded above by the logarithm of the number of bins. By construction, in a perfect bi-univocal relationship, the number of bins in one dimension is also the number of non-empty bins in the joint distribution, all sharing the same number of points. The higher the MI, the stronger the underestimate. On the other hand, due to the finiteness of the sample, the estimate has a positive bias (39) under independence of the variables or for small values. Indeed, any random sampling fluctuation will naturally result in a greater than zero estimate of the MI, even for independent variables. Formulae to correct for small sample sizes, such as Grasseberger correction (40), have been derived under the assumption of independent samples. However, such independence is violated by our equiquantal estimator as it imposes explicit bounds on the sums of rows and columns, rendering these corrections invalid — although they show relatively good agreement with our estimates for independent variables.

These biases have opposite signs and non-trivial tractability, and we thus addressed them numerically by evaluating the MI on samples from random bivariate Gaussian distributions with predetermined correlations (therefore known mutual information values according to Eq. (1)) and sample size S equal to the series length. Specifically, we calculated MI for 50000 bivariate random samples of size *S* for each correlation value ranging from 0 to .995 in increments of .005. The average of the 50000 MI estimates approximates the expected sample mutual information for each given mutual information value. This process yields a monotonous function (with linear approximation applied if needed to ensure monotonicity for correlations near zero) that relates the true mutual information to its expected numerical MI estimate. The inverse of this function then allows us to adjust each estimated MI to produce a more accurate bias-corrected estimate of the true mutual information.

We derive the maps assuming that the number of bins N and samples S and the true MI are the only determining factors for bias. Every combination of a number of bins and a number of samples requires a different map. This approximation holds for a wide range of observed relations spanning over 98% of our datasets’ connections.

Preliminary analyses with KNN and KDE estimators showed no substantial change at the price of much slower estimation. We explored in more detail the KNN estimator (41) (adapting the code from Tigramite (23)) and found that, while it’s superior to the binning approach in terms of separation of empirical and shadow datasets (thus in principle more sensitive to the removal of non-linearity) the estimates of RNL and the localisation of nonlinear connections stay essentially unchanged (see SI for more). Even if they account for non-independent samples, we excluded other MI estimators such as those including embedding in higher-dimensional spaces (42, 43), as they require even larger samples for a correct distribution estimate.

#### Non-linearity estimate

We define theoretically the non-linearity of a bivariate distribution through the extra-Gaussian information Eq. (2), i.e., the amount of information shared between two variables on top of that captured by their linear correlation. Indeed, of all distributions with Gaussian marginals, a multivariate Gaussian is maximally entropic for a given covariance matrix, i.e., has the minimum MI. Any deviation from Gaussianity will yield a higher MI.

Note that the above-mentioned assumption of Gaussian marginals does not imply a substantial loss of generality for quantifying dependence. In particular, Spearman’s correlation coefficient and MI are independent of the marginal distribution. Moreover, an approximate Gaussian distribution is often assumed, and every sample distribution can be mapped to Gaussian marginals with a monotonous transformation.

Indeed, we enforced Gaussian marginals via rank-normalization to ensure precise non-Gaussianity estimates; this is done by using the sample ranks to estimate the percentile *π*_*i*_ for each sample *x*_*i*_ and replacing the original value with the one corresponding to the same percentile in the standard normal distribution 𝒩 (0, 1).

We assess non-linearity’s contribution to the TMI, comparing it to the information conveyed only by linear correlations. For each dataset and subject, we calculated the MI on 99 random realizations of multivariate time series that preserve the linear structure but remove the nonlinear components. If the original time series has a Gaussian dependence structure, the original data MI would be similar to that of the surrogates, up to some random error due to the variability of the estimate and surrogates. Conversely, if the original data had substantially higher MI than the surrogates, this would indicate the presence of nonlinear dependencies.

We created the surrogates using the multivariate Fourier transform (FT) method (32, 44–46), which generates realizations of multivariate linear stochastic processes preserving the individual spectra and cross-spectra of the original time series. Specifically, each surrogate is obtained by adding identical random phases to corresponding frequency bins of the series’s FT while keeping the amplitude magnitudes unchanged. The series were then transformed back into the time domain using the inverse FT. These surrogates retain the dependence structure that a multivariate linear stochastic process can explain while destroying any non-linearity.

Comparing the MI estimate of the data and the “linear” surrogates, rather than directly using the linear correlation of the data, has two advantages. First, correlation and MI estimators have different characteristics regarding bias and variance, and surrogates thus allow for a direct quantitative comparison between nonlinear and linear connectivity. Second, surrogates represent a suitable null model for a direct statistical testing of differences. However, while these estimates are valuable for hypothesis testing— as we did to assess localization—we use the mean of these 99 values to quantify the relative difference. We refer to this as “Gaussian” MI, which closely approximates the MI of a bivariate Gaussian distribution. The Relative Non-Linearity (RNL) is the fraction of the TMI that exceeds the Gaussian MI. Note that, to obtain robust estimates of RNL, throughout the paper, it is reported at a global level, i.e., before taking the fraction, both TMI and Gaussian MI are first averaged over all pairs of signals.

As the correction map depends on the number of points used to estimate the MI, using oversampled (i.e., temporally correlated rather than independent) data would introduce some bias. The correction map reduces the measured value for low MI, which is typical for most connections. This necessary correction is smaller when the map is computed for a larger sample (i.e., lower bias). Using an oversampled sequence would lead to using a high-sample-count map, while the non-independent samples would still behave as if they were in a smaller number. As this applies to the original data and the surrogates, both measures would be overestimated by different amounts depending on their MI. This would convert to a negative bias in the fraction defining the RNL.

We acknowledge that for fMRI data, given the band-pass filter, the current sampling rate of 0.5 Hz may lead to oversampling. However, reducing the sampling rate to 0.18 Hz would give time series too short to get any reliable estimate of MI. In this case, the relative amount of nonlinear information thus has a slight negative bias. However, as this affects the shadow dataset similarly (see below), every result relative to it remains valid, as will the significance tests.

#### Shadow datasets

Following (14), we compared each result against a control analysis using linear “shadow” datasets. This approach allows us to account for any potential biases in the generation of surrogate distributions, such as those caused by small sample sizes. For each session, a shadow dataset was created as a multivariate FT surrogate of the original, marginally normalized dataset, thereby preserving only the original data’s linear structure (autocorrelation and cross-covariance).

We then applied the same processing to the original data and the shadow, including initial normalization, generation of multivariate surrogates, and computation of MI and RNL. This approach allowed us to replicate the entire procedure using a dataset with the same correlation and autocorrelation structure and purely linear interactions. This ensured that any potential bias in our findings due to the algorithm’s numerical properties was adequately controlled. We compared the RNL between the original and shadow datasets using a paired t-test. All group-level tests applied a family-wise corrected significance threshold of *p* = 0.05.

For Band-Limited Power (BLP) data, we considered two options for generating the shadow dataset. In the naïve version, we extract the power from each band and then surrogate the power values. This procedure destroys the sequence’s non-linearity and those that might have been introduced while computing the power. In the second version, we first generate a surrogate voltage sequence and then extract new power values from this.

### Amount of non-linearity

We relied on session-wise averages in each dataset to get the global fraction of non-linearity, RNL. In pure linear relationships, the total information measured on data should be indistinguishable from surrogates. We estimate the relative amount of nonlinear information in the brain—according to the different modalities—as the relative difference between the average of the total MI across all sequence pairs and the average of the Gaussian MI across all pairs and surrogates.

Despite individual and experimental fluctuation or any systematic bias in the estimate, session-wise relative differences for linear data should stay close to the estimate from the shadow dataset. A significant difference between empirical data and shadow datasets is the sign of the presence of non-linearity.

### Localisation of non-linearity

We evaluate the localization of non-linearity leveraging the statistics of the surrogates. If a region (or, equivalently, electrode) pair has linear interaction, repeated measures across sessions or subjects should give results analogous to the surrogates. We average the TMI and the 99 surrogate MIs across all subjects for each pair of regions, obtaining one average TMI and 99 average surrogates. We then use the mean and standard deviation of the surrogates to compute a *z*-score for the empirical TMI. We used Holm-Bonferroni (*α* = 0.01 to provide a reasonably conservative estimate) corrected *p*-values over empirical and shadow datasets together to assess which connections are significantly nonlinear across subjects. We use *z*-scores instead of the relative non-linearity computed on individual pairs, as the latter introduces a strong correlation between empirical and shadow datasets due to the shared variations in correlation.

Lastly, we treated the significant *z*-scores as weights in an undirected weighted graph and checked how the node degree compares with a random graph with the same link strength distribution. This comparison allows us to visualize regions with significantly higher or lower amounts of nonlinear connections than chance (here, *p <* 0.05 uncorrected to provide liberal yet spatially more informative visualization).

This analysis is possible only on the fMRI and EEG data, as other modalities do not comprehensively sample the whole brain. We also repeated the same steps on the fMRI data with raw preprocessing to highlight the role of artefact removal in the observed non-linearity.

### Reliablity

For any (nonlinear) FC measure to be potentially relevant as an individual characteristic (e.g., as a diagnostic biomarker), the estimator provided to the researcher must be reliable across sessions.

We compute the strength of the connectivity in three ways: Pearson’s correlation of the rank-normalized values, TMI, and the Gaussian MI estimate through Pearson’s correlation. The latter is obtained according to Eq. (1), with *r* being the sample Pearson correlation over the rank-normalized sequence. This estimate of information is faster to compute than TMI, but only accounts for linear relationships.

We also compared how well measures of FC through Eq. (1) or TMI in one session predict FC (estimated with MI) in a different session. We chose the AAL90 atlas for this task as it is closer to potential applications where reliability might be relevant.

We assume that high or low connectivity between specific regions is a marker of scientific or clinical interest. In this case, we desire that subjects ranking high or low in one session do so also in another. For each connection, we compute the Spearman correlation of the strength of connectivity of the subjects over two sessions. Note that we don’t linearise the matrices, but compute the correlation per connection and report averages across connections.

### Sources of non linearity

For EEG, iEEG, and single-unit spikes, we show examples of sources of non-linearity at the whole brain level. Firstly, we compute the RNL on BLP sequences for EEG, contrasting two ways of deriving the shadow dataset.

Secondly, in iEEG data, we show the evolution of RNL throughout a seizure. We used a sliding window of 45 s with 90% overlap for each frequency band. To provide more insight, the non-linearity is plotted along with the average power computed as the mean over all electrodes of the absolute value of the Hilbert transform averaged over each window.

Thirdly, we considered the single-unit spikes. We split the whole sequence into epochs of 15, 10, 6, 5, and 3 minutes, computed the RNL for each epoch, and then averaged over all sessions and epochs.

### Data

This work relies on five openly accessible datasets spanning four modalities: fMRI, EEG, iEEG, and single-unit spikes.

#### fMRI data

We used two different datasets of fMRI data: the public dataset used in (47)(ESO245) and the MPI-Leipzig Mind-Brain-Body dataset (48, 49) (LEMON).

#### ESO245

This dataset contains 10 minutes of resting-state eyes-closed functional magnetic resonance from 245 healthy subjects (148 right-handed, 132 females, mean age 29.22/standard deviation 6.99) acquired as healthy controls as part of the ESO project. Participants were informed about the experimental procedures and provided written informed consent. The local Ethics Committee of the Institute of Clinical and Experimental Medicine and the Psychiatric Center Prague approved the study design. The acquisition included T1-weighted and T2-weighted anatomical scans not used in this study. The scanner was a 3T MRI scanner (Siemens; Magnetom Trio) at the Institute of Clinical and Experimental Medicine in Prague, Czech Republic.

Functional images were obtained using T2-weighted echo-planar imaging (EPI) with BOLD contrast. GE-EPIs (TR/TE = 2,000/30 ms) comprised 35 axial slices—acquired continuously in descending order covering the entire cerebrum (48 × 64 voxels, voxel size = 3 × 3 × 3 mm^3^) (47).

#### Preprocessing

For the present study, we used the data already processed^∗^. We report the essential aspects of the processing while referring to (47) for the details.

The preprocessing followed the default pipeline of the CONN toolbox (McGovern Institute for Brain Research, MIT, USA) with 12 head motion parameters and five white matter components in the Component-based Correction (CompCor). The authors in (47) also detrended the resulting time series and applied band-pass filtering with cut-off frequencies 0.009-0.08 Hz. This pipeline is the “stringent” preprocessing compared to the “moderate” preprocessing (6 head motion parameters, one white matter component, cut-off frequencies 0.004-0.1 Hz) and “raw” (no CompCor nor filtering). The results in this paper use the “stringent” preprocessing unless explicitly stated otherwise.

#### Choice of the atlas

We used two sets of atlases available in (47). The first includes 23 different parcellations using the Craddock atlas and a number of ROIs between 10 and 950 to investigate the effect of spatial averaging and region size on non-linearity. The second is the widely used AAL atlas with 90 regions to assess non-linearity localization.

The normalized cut spectral clustering used in Craddock parcellation can yield a number of ROIs smaller than desired—i.e., empty clusters—and some ROIs can fall outside the GM mask for some subjects. This effect is more pronounced with decreasing region size and is observed in this dataset for all sizes larger than 200. The most significant number of ROIs is 840 against a seed of 950. The average number of voxels included in each ROI of the Craddock atlas thus varies from almost 2 × 10^3^ with ten regions to about 20 voxels with 840.

For each set number of ROIs in the Craddock atlas, we removed the ROIs that were empty for any subject from all subjects. We discarded the three subjects with the most empty regions to avoid discarding too many ROIs. The resulting dataset for region-size effect analysis includes 242 subjects with parcellation in 10 to 691 regions.

#### LEMON

We included a subset of the MPI-Leipzig Mind-Brain-Body dataset (48, 49) to assess non-linearity test-retest reliability^†^. The subset contains the 14 subjects (1 female, ages 20 to 35, reported in 5-year bins, mode 25-30) with at least three resting state measurement sessions, all with the same TE (to avoid possible effects due to scanning parameters). We applied the same CONN toolbox default, “stringent” preprocessing pipeline and chose the AAL 90 parcellation.

#### EEG data

The EEG data for this study derives from 8 minutes of resting-state eyes-closed recording from 215 healthy participants in the Max Planck Institut Leipzig Mind-Brain-Body Dataset – LEMON. We report here the main features of the data referring to the dataset presentation papers (48, 49).

#### Data acquisition

The EEG data was acquired with a sampling frequency of 2500 Hz using 62 channels (ActiCAP, Brain Products GmbH, Gilching, Germany) according to the 10-10 system with one VEOG electrode. The total acquisition lasted for 16 minutes, divided into 60 s blocks with eight eyes closed blocks interleaved with eight eyes open blocks.

#### Data preprocessing

We downloaded the preprocessed version of the dataset containing 204 subjects^‡^. The preprocessing included band-pass filtering (1–45 Hz), downsampling to 250 Hz, removal of corrupted channels, and eye movement and heartbeat removal via ICA. We excluded from the analysis seven electrodes frequently missing (T7, T8, Cz, F7, CP6, PO10, Fp2) and considered the 150 subjects with all the remaining 54 channels available.

We further processed the data in Python using the MNE (50) and SciPy (51) packages. At this stage, the sequences are split into segments up to 60 s long. We obtained the five usual frequency bands (*δ* = [1, 4] Hz, *θ* = [4, 8] Hz, *α* = [8, 12] Hz, *β* = [12, 30] Hz, *γ* = [30, 44] Hz) from each segment using an IIR Butterworth filter of order 4 in two forward and backwards passes to minimize phase distortion. We obtained the scalp voltage by down-sampling to 1.25 times the Nyquist frequency of the upper limit of each band of the filtered sequence to avoid biases (see “MI estimator”).

We derived the scalp BLP from the filtered signal by applying the Hilbert transformation and taking block averages of the modulus over 125 ms windows. We obtained the naïve shadow dataset for BLP sequences, treating the power time series as the source data, i.e., computing multivariate FT surrogates after applying the Hilbert transformation and the averaging right before RNL estimation. Instead, for the non-naïve shadow dataset, we first compute the surrogate and then extract the power. Finally, we obtained the three epochs used for RNL and reliability estimation by chaining together segments for a total of 124 s.

#### iEEG data

The iEEG dataset is derived from the open dataset published by the Sleep-Wake-Epilepsy-Center (SWEC) of the University Department of Neurology at the Inselspital Bern and the Integrated Systems Laboratory of the ETH Zurich (52). The original dataset^§^, contains 2656 hours of anonymized and continuous intracranial electroencephalography (iEEG) of 18 patients with pharmaco-resistant epilepsies. All the patients gave written informed consent that their iEEG data might be used for research and teaching purposes. Given the extensive size of this dataset, for each subject, we selected the central 124 s of the first hour that was at least 45 minutes away from any seizure based on metadata and manual inspection.

#### Data acquisition

The iEEG signals were recorded intracranially by strip, grid, and depth electrodes. The recorded signal was saved at a rate of 512 or 1024 Hz after band-passing between 0.5 and 120 Hz with a double-pass fourth-order Butterworth filter. An epileptologist visually inspected all the recordings, marked each seizure’s onset and end, and removed corrupted channels.

#### Data preprocessing

The 18 subjects have between 24 and 128 electrodes of undisclosed type in undisclosed locations. Also, it is not reported which electrodes are located in the epileptic foci. To improve data homogeneity, we randomly selected 24 electrodes for each subject. We band-passed and down-sampled the time series as we did for the EEG electrode voltage and selected a segment of 124 s. We extracted three additional subsets of the data. In the first, the sequences have the same starting time but cover different time lengths (up to 22 minutes and 44 s) depending on the frequency band to allow in each band the same number of samples as for band gamma with 124 s. The other two correspond to windows of 124 s extracted 24 and 48 hours after the first one. In case of a seizure or a total length of the recording under 48 hours, we selected seizure-free hours, trying to keep the distance between windows as close as possible to 24 hours.

#### Single Unit Spikes data

The last dataset we include is derived from the Allen Brain Observatory – Neuropixels Visual Coding dataset (53). The dataset^¶^ contains individual units isolated in 58 mice from six probes inserted in the visual areas. The dataset section relevant to this work includes spiking times from individual units and metadata about unit localization and quality. We considered the 26 specimens that underwent the “functional connectivity” experiment. This experiment included 30 minutes of spontaneous activity, i.e., resting-state data.

#### Data preprocessing

With this dataset, we aimed to explore the effects of averaging on non-linearity with the finest granularity. We first selected units with reasonable quality measures (expected fraction of missed spikes ≤ 0.08, maximum inter-spike interval violations = 0.2 (54)). Then, we created groups of 32 units from the same brain structure. If a structure had enough units to form more than one group, we determined the groups so that the units were spatially separated. When the number of exceeding units is not enough to create an extra group, we discarded those of lower quality weighting in order: ISI violations, quality of isolation from other units, and the expected fraction of missed spikes. We kept only the 16 sessions that allowed the identification of at least nine groups.

In this case, our data will be the instantaneous firing rate estimated at regular 60 ms intervals. The instantaneous firing rate is estimated as the inverse of the weighted local average of inter-spike intervals. Given the series of spiking times 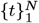 we define the instantaneous firing rate at time *τ* as *fr*(*τ* ) : = 2 · [(1 − *α*)(*t*_*i*_ − *t*_*i*−1_) + (*t*_*i*+1_ − *t*_*i*_) + *α*(*t*_*i*+2_ − *t*_*i*+1_)]^−1^. Where the spiking time *t*_*i*_ | *t*_*i*_ < *τ* ≤ *t*_*i*+1_, and 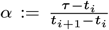. With this definition, the instantaneous firing rate is a continuous function of time and equals the inverse of the average of neighbouring spiking intervals when evaluated at spiking times. For every session, we derived 6 sets of time series, taking the average firing rate of 1 to 32 units (doubling the number at each step), assessing thus the effect of spatial averaging.

## Code availability

The code and scripts for the full reproduction of these results and the estimation of the nonlinear content of timeseries is available at: https://github.com/cobragroup/nonLinearity.

## ACKNOWLEDGMENTS

The publication was supported by ERDF-Project Brain dynamics, No. CZ.02.01.01/00/22 008/0004643, the Czech Science Foundation projects No. 21-32608S and No. 21-17211S.

## Footnotes

∗ Available at doi.org/10.17632/crx7d22pym.4

¶ Available at: https://allensdk.readthedocs.io/en/latest/visual_coding_neuropixels.html

† Available at: https://fcon_1000.projects.nitrc.org/indi/retro/MPI_LEMON/downloads/download_MRI.html

‡ Available at: https://fcon_1000.projects.nitrc.org/indi/retro/MPI_LEMON/downloads/download_EEG.html

§ Available at: http://ieeg-swez.ethz.ch/

## Notes

http://doi.org/10.17632/crx7d22pym.4

https://fcon_1000.projects.nitrc.org/indi/retro/MPI_LEMON/downloads/download_MRI.html

https://fcon_1000.projects.nitrc.org/indi/retro/MPI_LEMON/downloads/download_EEG.html

http://ieeg-swez.ethz.ch/

https://allensdk.readthedocs.io/en/latest/visual_coding_neuropixels.html

### Competing Interest Statement

The authors have declared no competing interest.

### Summary of Updates

Introduction of additional analysis, new comparison of our results with multiple alternative measures as recommended by the reviewers, clarifications in the methods, and improvements in the general discussion.

