## Supplemental Information for "Nonlinear brain connectivity from neurons to networks: quantification, sources and localization"

**This PDF file includes:**

Supporting text
Figs. S1 to S6
Tables S1 to S2
SI References

### Supporting Information Text

**Dependence of the shadow dataset RNL on the sequence length.** In the main text, a trend was observed in the RNL measured in the shadow dataset when varying the frequency band or the width of the time window for the average spiking rate in mice. This behavior shouldn't come as a surprise. Using fewer samples—as is the case of low-frequency bands and spiking rates averages over large windows—the fluctuations in the estimated MI are more significant even for the linear surrogates, including those forming the shadow dataset.

The MI can't be lower than zero, so its distribution is right-skewed. This effect of the lower bound is reduced for higher information values (equivalent to correlation for the surrogates), but most of the correlation values observed are small. This introduces, on average, a positive bias in the estimates of RNL on the shadow dataset. Indeed, values exceeding the surrogates' average are more likely to be further from the average than values smaller than the average (which are strictly bound by zero from below).

To further validate this observation, we repeated the analysis for the iEEG on sequences with the same number of samples for each frequency band. Given our resampling policy to capture the typical temporal scale, this implies that if for band $\gamma$ we keep 124 s, to get an equal amount of samples in the band $\delta$, we need over 22 minutes of recording. The first 124 s are the same as used in the main text. The effect can be appreciated in Figure S1 compared to Fig. 1C of the main manuscript. The RNL for the shadow datasets becomes centered around zero for all bands, excluding a bias for lower frequencies. At the same time, the RNL for the empirical dataset seems to grow in lower bands. This growth is most likely due to non-stationarities of brain activity over the extended time of the recording of the natural (non–resting-state) activity of the subjects.

### Contribution of epileptic activity to RNL

The iEEG data comes from epileptic subjects during pre-surgical assessment. This implies that, even away from seizure, the brain may undergo some epileptic activity, mainly in the form of Interictal Epileptic Discharges (IED) and High-Frequency Oscillations. This kind of activity usually involves more than one electrode, and the connectivity thus acquires a (nonlinear) contribution due to the different amplitude of the spikes compared to regular activity. This is a source of apparent non-linearity of the dependence and increased FC. Researchers may consider whether such dependence due to sparse outlying events should be regarded as proper functional connectivity of interest or an artifact of the outliers. In any case, the presence of IEDs might be better addressed with specific tools than lumped into an overall functional connectivity estimate together with the spontaneous healthy activity.

We implemented the algorithm described in (1) to identify and count the number of spikes occurring on all electrodes during the 124 s used to estimate RNL in the main text. In Figure S2, we show the relationship between the number of IEDs and the observed RNL in each band for each subject. While there's a gradual increase in the slope of the relationship, only bands $\beta$ and $\gamma$ show a significant correlation capable of explaining most of the observed RNL. This result can be linked to the overlap between these two bands and the band of frequencies more impacted by IEDs. A similar analysis on individual electrode pairs yields analogous results with correlation for lower frequency bands that are significant but reduced.

Overall, accounting for IEDs reduces the amount of unexplained non-linearity that might also be observable in healthy brains. However, this modality still confirms a more considerable relevance of non-linearities in iEEG than fMRI and EEG, especially in lower frequency bands.

### Localisation of non-linearities in fMRI data

We used the node degree in a network weighted by the significance of connection non-linearity to determine which regions—if any—had an increased concentration of non-linearity. However, in the case of fMRI, there is a substantial correlation between the regions with more significant non-linearity in the empirical and shadow datasets.

We suggest that most regionally-specific non-linearity is due to artifacts and is removed during preprocessing. Indeed, when using stringent preprocessing, the correlation between region degrees is 0.37 ($p = 0.00028$, uncorrected). The correlation grows to 0.58 with moderate preprocessing and 0.69 with the raw one. In Figure S3, we report the adjacency matrices for empirical and shadow data with the three preprocessing pipelines. While the correlation is based on the sum over rows of the connection strengths, we notice similarities in the structure of the matrices, especially in the occipital and central areas.

### Test-retest reliability of EEG and iEEG results

For EEG and iEEG data, we don't have separate sessions available (as we had for fMRI). However, we approximate the effect of a test-retest experiment by taking separate segments of the same session. This practice would provide an optimistic view of the reliability as electrode position and characteristics will be precisely the same, and any slow variation in brain activity is excluded. For the iEEG data, we selected windows 24 hours apart to get a more realistic estimate while choosing the same time of the day to suppress diurnal cycles.

In Table S1, we report the average over all the electrode pairs of the reliability (assessed by Spearman correlation, across subjects, of functional connectivity in the two sessions) of functional connectivity estimated via TMI or correlation. When predicting TMI in a different session from the correlation $r$, we transform the correlation to the analytical MI estimate given by $MI = -\frac{1}{2}\log(1 - r^2)$.

As expected, the reliability is higher for electrophysiological signals than in the fMRI case, particularly for EEG. In this case, all three "sessions" happen within 15 minutes with exactly the same electrode positioning. We also note that there is only one case (EEG, $\delta$-band) where the prediction of TMI from correlation is better than from TMI. However, the difference is less than 1% in most cases, and correlation reliability alone is always higher. This suggests that noise, especially in low-frequency bands, negatively impacts the TMI measures more than the correlation.

### Alternative measures of Mutual Information

The binning method for the estimation of MI is fast, but the quality of its estimates has been surpassed over the years by more modern, yet computationally more demanding, approaches. We thus repeated part of our analysis using the KNN estimator (2).

The KNN estimator has a lower estimation bias than the binning approach, making its numerical correction unnecessary. This comes with a cost in computational complexity. The binning algorithm to sort once the $n$ points of all $s$ time series ($o(sn \log n)$) and count for every pair of series the occupancy of every bin ($o(s^2 n)$). Conversely, the KNN—even when implemented with fast kd-trees—requires for every pair of series to build a tree (at least once) and then, for every sample, to find the radius of the $k$-th neighbour and search that span. This last operation gives the leading scaling factor of $o(s^2 n^{\frac{3}{2}})$, and it's performed twice for each node with the search of the $k$-th neighbour scaling only slightly better. In practice, the non-leading terms and the constants strongly impact the running time.

The first observation is that using KNN allows for a better separation between the empirical and shadow datasets. A simple threshold that separates the two datasets based on RNL has an AUC of .96 when using KNN and .75 when using binning. However, this improvement doesn't come with an equally large difference in the measures of RNL and the localisation.

In figure S4, we present the impact on RNL and localisation of the different algorithms. For fMRI, where the short sequences lead to less manageable bias, we observe a clear difference between the binning and KNN approaches. The large values with 200 neighbours are likely related to the large bias of the estimator in this case (indeed, large $k$ is advised not to measure MI but rather to test the presence of dependency, thanks to the small bias when MI is zero and a low variance (2)). The following results are computed with $k = 7$ to keep low bias and avoid the high fluctuations of very small $k$ (2, 3). For the KNN estimator, the imposition of normal marginals is reported to reduce the estimation bias (3). In the case of EEG, the values of RNL are very similar with the two approaches, as are the maps highlighting the most nonlinear regions, which show some qualitative difference only in band $\beta$. The localisation analysis on fMRI data shows the same significant correlation between empirical and shadow datasets, which increases with reduced preprocessing.

### Alternative measures of dependence

While Mutual Information is the principled measure with a theoretically-grounded minimum for Gaussian data, we can derive other estimates of the relevance of non-linearity, e.g., by comparing the connectivity matrices obtained on the empirical data and those on the shadow dataset.

To check the impact of non-linearity on different measures, we computed connectivity matrices on the empirical data. We recorded the average $L_2$ norm of the difference with the matrices from surrogates. We then repeated the procedure on the shadow dataset. If the connectivity measure is sensitive to non-linearity, the difference between empirical and surrogates should be, on average, larger than between shadow and surrogate.

We quantify the sensitivity to non-linearity with the AUC of a threshold separating empirical and surrogate data, and consider the following measures:

1. Effective Connectivity at lag 1,
2. Chatterjee correlation (4),
3. distance-transformed Chatterjee correlation (5) considering pairs of samples with lags 1, 3, and 10.

Despite the ability of these measures to leverage directed relationships and some degree of state space information, their ability to differentiate empirical data from their linear transformation is at best comparable to MI, as shown in table S2.

We did not compute the AUC for all pairs of data and measures due to the high computational cost of these measures. Effective Connectivity estimate by Conditional Mutual Information (which we compute using KNN to get better results) is significantly more expensive than KNN MI due to the additional dimension requiring more trees and more searches. Chatterjee correlation has a complexity $o(s^2 n \log n)$ due to sorting each sequence with respect to every other to get the ranks. Due to the transformation that increases the number of effective samples as the number of possible pairs of samples, the distance-transformed Chatterjee correlation has complexity $o(s^2 n^2 \log n)$. For this reason, we computed it only for the shorter series.

### Classification task

As discussed in the main text, more advanced and data-hungry approaches impose a tradeoff between expressivity and statistical power. We exemplified how this can have a practical impact in research practice by replicating part of the study described in (6).

Out of the five datasets used in (6), we chose the COBRE (Center for Biomedical Research Excellence*) dataset comprising rest-fMRI data to study schizophrenia and bipolar disorder (7). The task is to predict subjects with a schizophrenia diagnosis versus normal controls.

* cobre.mrn.org., https://fcon_1000.projects.nitrc.org/indi/retro/cobre.html

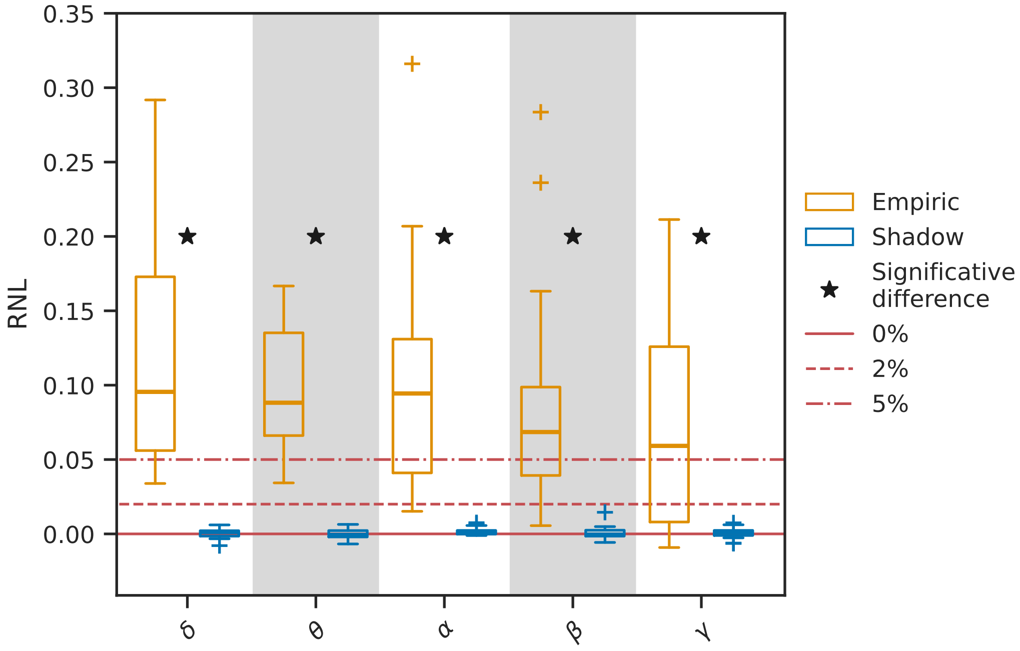


**Fig. S1.** Distribution over subjects of the Relative amount of Non-Linearity (RNL) for iEEG voltage using the same number of samples (S=12276) for all bands. The shadow dataset is a linear surrogate of the empirical one; see text for details.

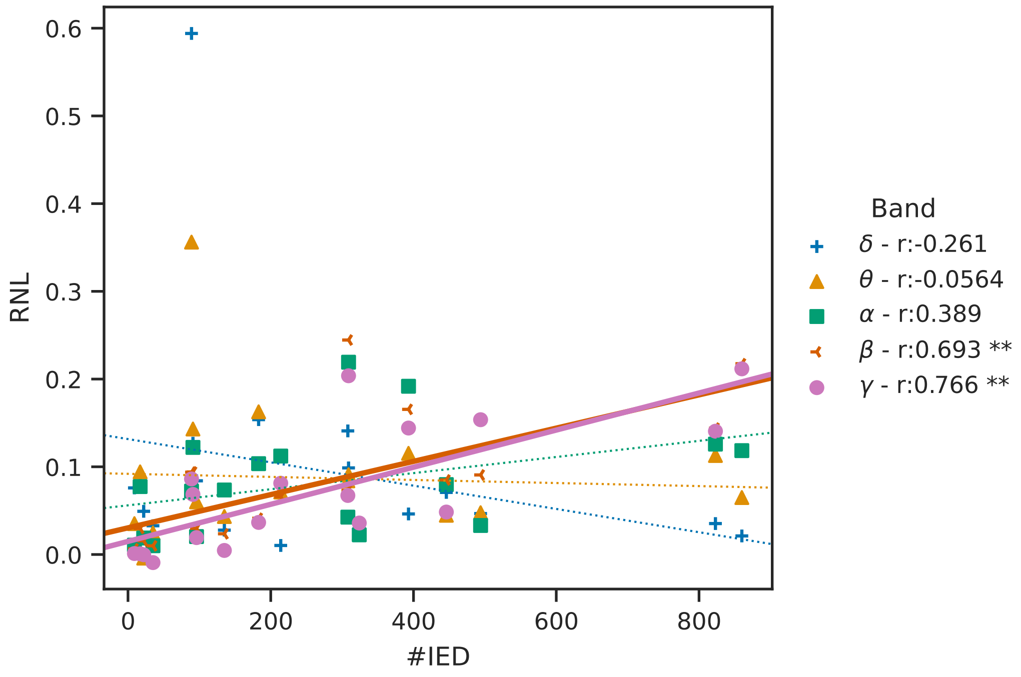


**Fig. S2.** Relationship between number of IEDs and RNL in different bands. ** inficates $p$-value $< 0.01$, Bonferroni corrected.

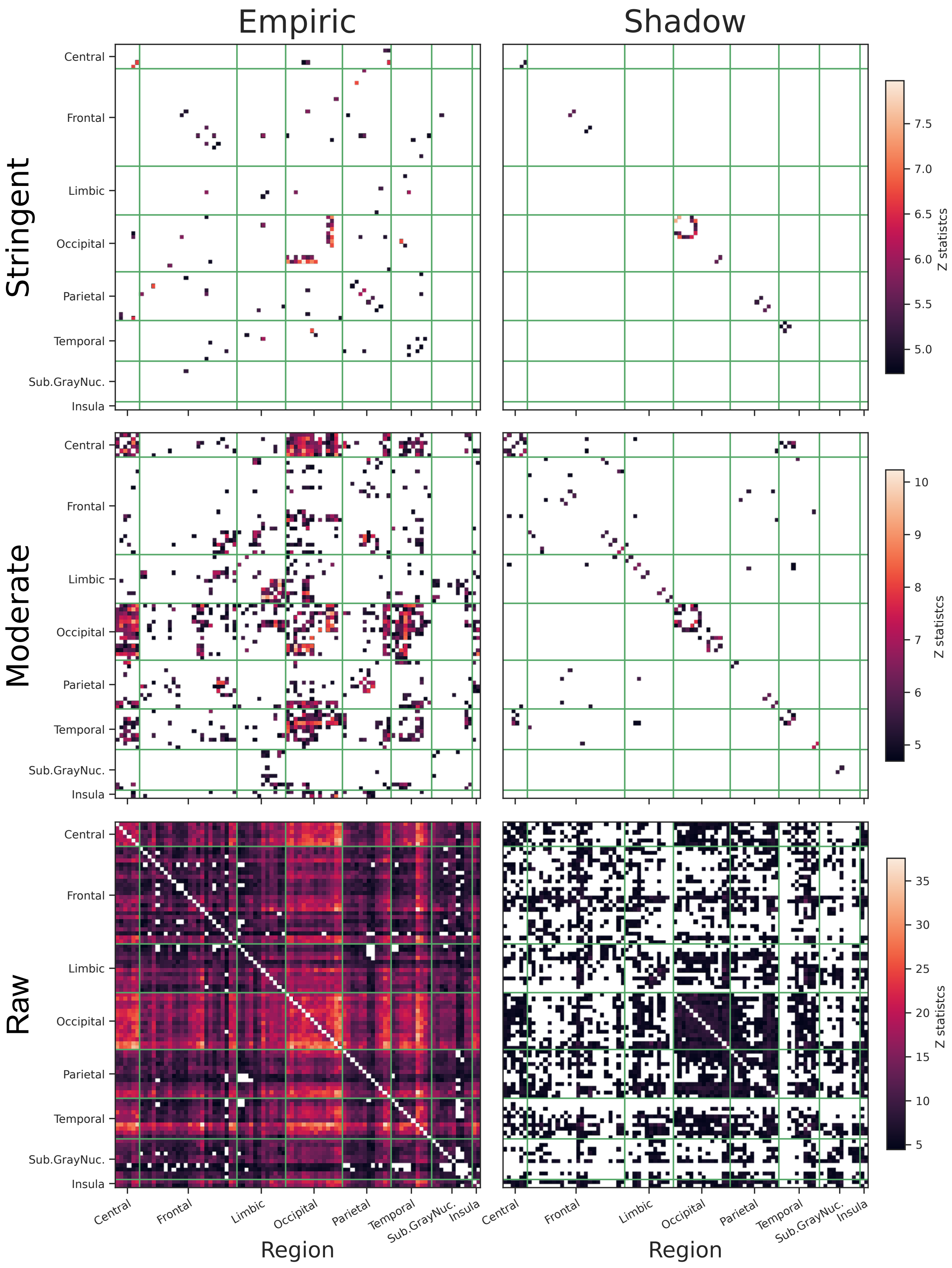


**Fig. S3.** Maps of the significant z-scores of TMI normalized according to the surrogate distribution.

**Table S1. Reliability and cross-predictability of functional connectivity. Average over connections of the Spearman correlation of functional connectivity estimates between sessions. For each session pair, the ranks can be determined using different estimators. $I_{\text{Gauss}}$ is the Gaussian approximation to Mutual Information, where the MI is estimated as a function of Pearson's correlation (according to Eq.(1) in the main text).**

| Modality | Band | TMI | $I_{\text{Gauss}}$ predicting | | Pearson |
|---|---|---|---|---|---|
| | | TMI | TMI | $I_{\text{Gauss}}$ | Pearson |
| fMRI | — | .134 | .142 | .153 | .277 |
| EEG | $\delta$ | .852 | .861 | .878 | .896 |
| | $\theta$ | .905 | .899 | .909 | .921 |
| | $\alpha$ | .873 | .860 | .878 | .911 |
| | $\beta$ | .904 | .900 | .913 | .930 |
| | $\gamma$ | .863 | .861 | .862 | .885 |
| iEEG | $\delta$ | .715 | .715 | .752 | .844 |
| | $\theta$ | .711 | .704 | .731 | .823 |
| | $\alpha$ | .699 | .675 | .685 | .733 |
| | $\beta$ | .740 | .664 | .691 | .809 |
| | $\gamma$ | .792 | .772 | .787 | .835 |

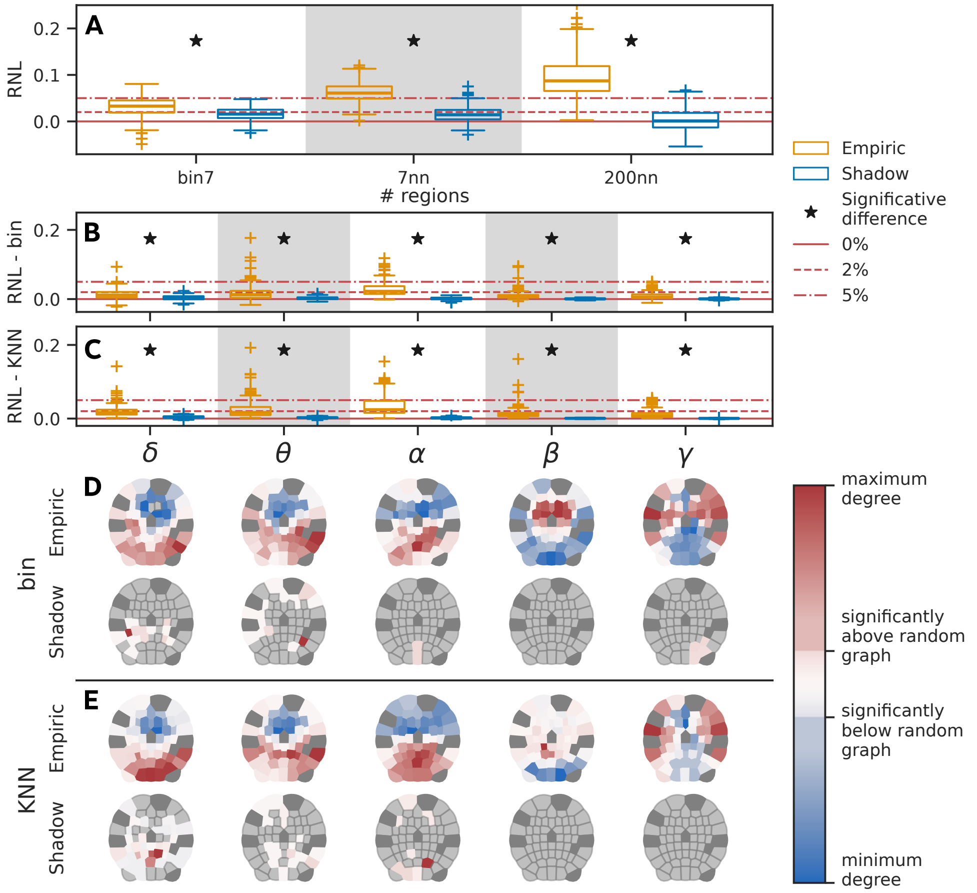


**Fig. S4.** Comparison of the RNL using binning estimate in (**A**) fMRI with AAL atlas and (**B**-**C**) EEG. **D**-**E**: localisation of non-linearity in EEG using binning or KNN with 7 neighbours. Panels **B** and **D** are the same as panel **B** in Figure 1 and Figure 3 of the main text, respectively.

**Table S2. AUC of a simple classifier based on the average $L_2$ distance between data and surrogates for different connectivity measures. For the measures based on Chatterjee correlation, we also report the results on the raw preprocessing for fMRI, which exhibits larger apparent non-linearity (see localisation analysis). We don't report results where the poor scaling with the number of samples (faster than $o(s^2 n \log n)$) made the computation greatly impractical with longer series.**

| Data | Subset | MI | EC | $Chatt$ | $d^{(1)}_{Chatt}$ | $d^{(3)}_{Chatt}$ | $d^{(10)}_{Chatt}$ |
|---|---|---|---|---|---|---|---|
| fMRI | stingent | .87 | .86 | .61 | .54 | .58 | .58 |
| | raw | — | — | .76 | .62 | .63 | .66 |
| EEG | $\delta$ | .77 | — | .67 | .73 | .67 | .66 |
| | $\theta$ | .85 | — | .74 | .71 | — | — |
| | $\alpha$ | .94 | — | .85 | — | — | — |
| | $\beta$ | .93 | — | .74 | — | — | — |
| | $\gamma$ | .92 | — | .85 | — | — | — |
| iEEG | $\delta$ | .80 | — | .67 | .76 | .70 | .68 |
| | $\theta$ | .88 | — | .59 | .76 | — | — |
| | $\alpha$ | .85 | — | .66 | — | — | — |
| | $\beta$ | .87 | — | .65 | — | — | — |
| | $\gamma$ | .83 | — | .75 | — | — | — |

We used the AAL atlas despite having a marginally lower performance (5-10% loss compared to the best choice) for consistency with the rest of this work and because it does not require retraining for every validation fold. We chose a Logistic regression classifier with $\ell_2$ as this is the best performing classifier overall, particularly for COBRE.

For the connectivity parameterisation—where the authors in (6) considered partial correlation, correlation, and tangent space—we considered three groups of options:

1. marginal independent: Mutual Information and Spearman's correlation,
2. on native marginals: Pearson's correlation, MI on shadow dataset,
3. on normalised marginals: Pearson's correlation, MI on surrogate.

These groups always include one measure, potentially sensitive to non-linearity, and one linear measure.

In figure S5, we show the distribution over the cross-validation folds of the AUC of the classifiers trained with different connectivity parameterisations. The results are very similar for all six measures, with the only significant difference being that the AUC with Mutual Information is lower than with MI on surrogates. This again supports our claim that, on fMRI data, most of the information is in the linear part and linear measures offer the best performance.

The median AUC using Pearson's correlation on the original marginal is 0.753. These figures are consistent with the results in (6), accounting for the reduction in performance compared to the best choice of atlas and connectivity parameterization.

### Linearising effect of averaging

We show here an example of how a system with nonlinear interactions can have an increasingly Gaussian copula as a result of averaging. Without claiming this to be a realistic model, we considered two populations of $N = 1000$ Kuramoto oscillators. Each oscillator can be seen to loosely represent one cortical column, such that the signal from one node can be paralleled to the iEEG signal, and the signal from a thousand mimics the EEG.

We added sparse connectivity inside each population ($p_{in} = 0.12$ of two nodes being connected) and sparser connectivity between populations ($p_{out} = 0.08$), but with a stronger connectivity between "contralateral" nodes ($P(n_i^1 \leftrightarrow n_{i+l}^2) = 0.58$, with $l \in \{0, 31\}$, $i = 1, \ldots, N$). Each node has a spontaneous frequency $\omega_i \sim \mathcal{N}(\infty, \prime.\in)$. The dynamics is driven by a connectivity constant $K = 30$, Gaussian noise on the individual nodes $\varepsilon_i \sim \mathcal{N}(0, 0.2)$ and a common driver for each region $\varepsilon_r \sim \mathcal{N}(0, 3.2)$.

Suppose that the observable signal corresponds to the sine of the phase of the nodes in the absence of noise. With a lower value of $K$, the nodes are not synchronised and independent of each other $\left(r\left(\sin(\phi_i^1), sin(\phi_i^2)\right) \approx 0\right)$. With our choice of $K$, the nodes are locked in phase, giving an elliptical dependency when looking at individual nodes and almost perfect correlation when looking at averages of large enough groups of nodes. Indeed, the Kuramoto order parameter is close to $R \sim 0.872$ and the nodes group around the same average phase.

If we now add noise, we allow the system to explore more configurations. The median Kuramoto order parameter is still $R = 0.735$, but random fluctuations can bring it close to zero for brief moments. We can thus observe in Fig. S6 how the Gaussian-marginalised distribution of samples starts diamond-shaped when looking at a single node per region and approaches a Gaussian when larger groups of nodes are taken into account.

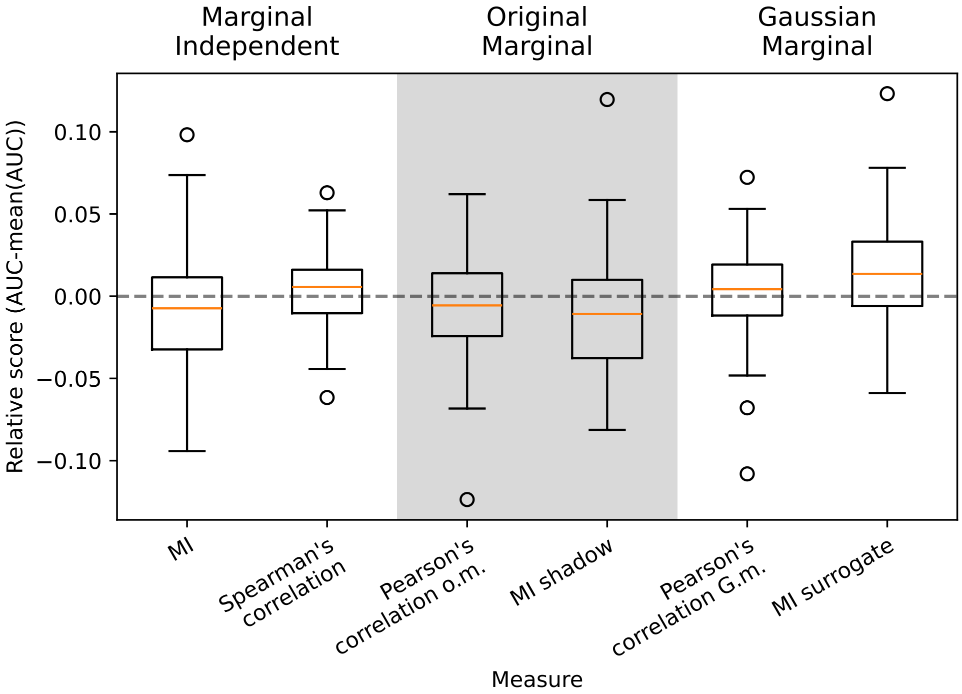


**Fig. S5.** Effect of the different measures on the AUC of a logistic regression classifier identifying subjects with a schizophrenia diagnosis versus normal controls over 100 cross-validation folds. To compensate for differences among folds, the mean AUC across measures is subtracted from the AUC of each measure for each fold as in (6).

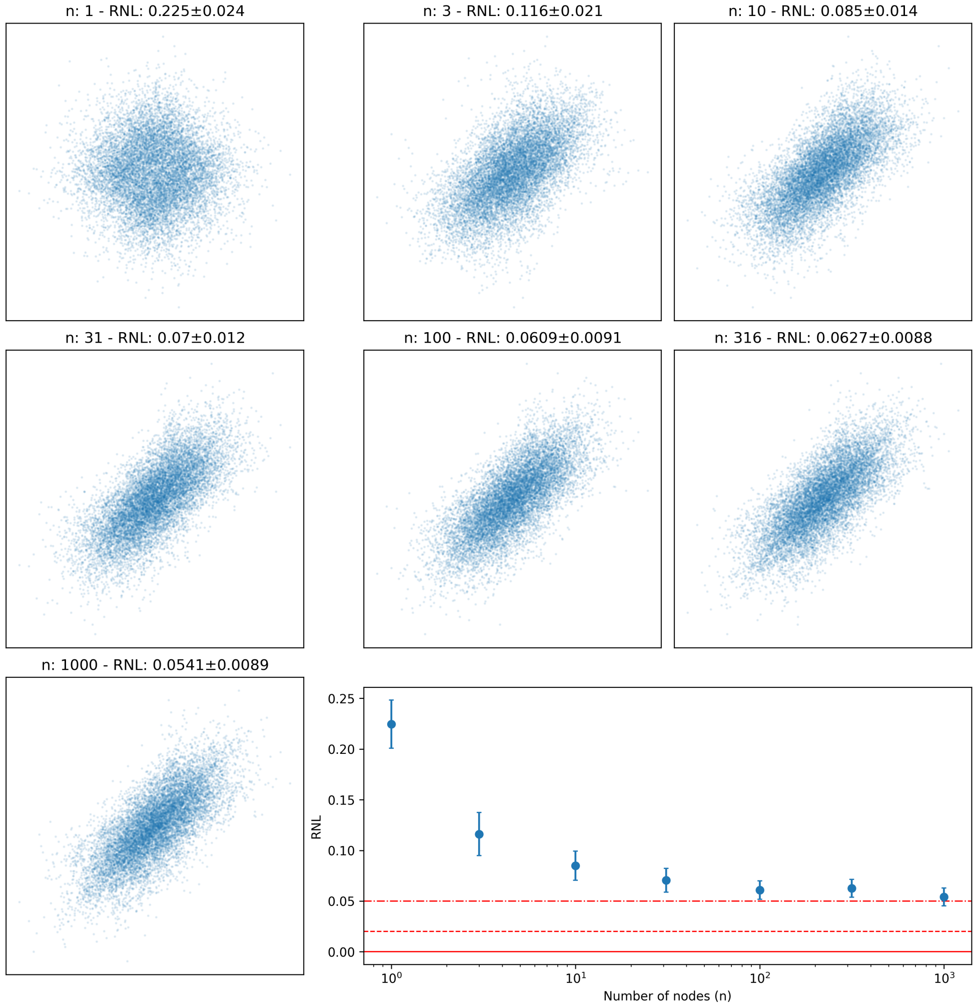


**Fig. S6.** Simulating the effect of spatial averaging on the RNL –distribution of samples and RNL in a network of Kuramoto oscillators. The scatterplots represent one sample distribution for each number of nodes. For each group size $n$ we obtain the sequences as $x^r(t) = \frac{1}{n}\sum_{i=1}^{n} \sin\left(phi_i^r(t)\right)$, with $r = 1, 2$ the region index, and we report them rank-normalised. The RNL is estimated as the relative difference of the average MI of $\frac{N}{n}$ independent groups of nodes and the average of their surrogates. The error is estimated from the spread of the MI of simulated data and surrogates.

 
